# Autoinsertion (LAiR): rapid functional reconstitution of integral membrane proteins into lipid bilayers

**DOI:** 10.1101/2022.09.21.508839

**Authors:** Albert Godoy-Hernandez, Amer H. Asseri, Aiden J. Purugganan, Chimari Jiko, Carol de Ram, Holger Lill, Martin Pabst, Kaoru Mitsuoka, Christoph Gerle, Dirk Bald, Duncan G. G. McMillan

## Abstract

Functional investigation of purified integral membrane proteins (IMPs) is hampered by the need to insert these hydrophobic proteins from the detergent-solubilized state into liposomal membranes. Here we report reintegration of IMPs into a lipid environment within minutes, an order of magnitude faster than currently used standard techniques. The new approach yielded optimal results for IMPs solubilized in the detergent lauryl-maltose neopentyl glycol (LMNG) and is therefore termed **L**MNG **A**uto-**i**nsertion **R**eintegration (LAiR). LAiR displays superior performance to standard methods in terms of protein activity, long-term stability and proton tightness of proteoliposomes. LAiR reconstituted vectorial control of membrane-bound activity by the transmembrane ion motive force, a property particularly important in mitochondrial function, which was undetectable by standard reintegration methods. LAiR also preserved fragile IMP properties that are prone to disruption upon reintegration, including long-term multi-subunit integrity, inhibitor susceptibility, and higher-order oligomeric states. LAiR proved suitable for reintegration into liposomes as well as into surface-tethered membrane bilayers, and was compatible with IMPs and lipids from prokaryotic and eukaryotic sources. We anticipate a broad scope for LAiR as a powerful tool in fundamental research, pharmaceutical applications, and biotechnology.

## Introduction

Membrane-embedded proteins fulfill a broad range of essential tasks in all living organisms, including cellular signaling and recognition, energy conversion, and transport of ions and metabolites. In line with these key biological functions, integral membrane proteins (IMPs) are highly represented among the targets of clinically utilized pharmacophores^1,2^. Yet, even after decades of efforts multi-subunit IMPs still remain extremely challenging to study.

Functional investigation of IMPs typically requires purification of the protein from a complex mixture, thereby avoiding interfering signals from other membrane components. For IMP purification, detergents are commonly utilized, and are viewed as simply a means of extracting the proteins from biological membranes and to retain the IMP solubilized in an aqueous buffer solution. To maintain the lipid environment, which may be needed for optimal function of a solubilized IMP^3^, amphiphilic polymers (amphipols) or lipid bilayers encircled by amphiphilic protein scaffolds (nanodiscs) can be utilized^4,5^. However, vectorial IMP functions cannot be assessed in the detergent-solubilized or nanodisc-solubilized state. Vectorial properties, such as transmembrane signaling, maintenance of ion gradients, and transport of ions or metabolites, are key elements of physiological IMP function.

Reintegration of purified membrane proteins into liposomes is a powerful tool to reconstitute and assess vectorial IMP properties and therefore widely utilized for functional studies. Reintegration of a nano-disc solubilized IMP into liposomes has been reported^6^, but the fate of the protein scaffold after reintegration is unknown and the scope of this utility needs to be explored. To facilitate incorporation of detergent-solubilized IMPs into the liposome, various approaches have been applied, including dialysis, gel filtration chromatography, rapid dilution, and adsorption of the detergent on the surface of polystyrene beads^7,8^ or combinations of these approaches^9^. Biobeads^®^ SM^2^ resin (Bio-Rad, referred to as “biobeads” hereafter), a nonpolar polystyrene adsorbent, have been applied for liposome reintegration of a broad variety of IMPs, including bacterial transport proteins^10^, ATPases from prokaryotic and eukaryotic sources^11,12^, plant photosystems^13,14^ and viral membrane proteins^15^. This method is exceptionally efficient for removal of detergents with a low critical micelle concentration, such as Triton X-100 and β-dodecyl-maltoside (DDM)^8^. DDM presently is the most widely used detergent for purification and crystallization of IMPs^16^ and liposome reintegration of DDM-solubilized IMPs by biobeads may be regarded as a “gold-standard” in the field. However, even in successful cases the resulting proteoliposomes suffer from proton leakiness and instability, preventing accurate measurement of vectorial functions. Furthermore, the incompatibility of very mild detergents (e.g. digitonin), imperative for retaining higher-order membrane protein assemblies, renders the *in vitro* study of these complexes in membranes to be out of reach.

In the current report, we present a new approach for efficient reintegration of IMPs into liposomes within several minutes, without the need for adsorbent beads. When applied on integral membrane proteins solubilized in lauryl maltose neopentyl glycol (LMNG), the enzymatic activity of the reintegrated proteins and the proton tightness of the proteoliposomes clearly exceeded the biobead standard over extended periods. This new approach, termed **L**MNG **A**uto-**i**nsertion **R**eintegration (LAiR), reconstituted the impact of a natural proton motive force on IMP activity, which was not detectable with the biobead method. LAiR also preserved delicate IMP properties such as the functional integrity and higher-order oligomeric state of fragile multi-subunit IMPs. We demonstrated that LAiR was highly compatible with both preformed liposomes and membrane bilayers tethered on an electrode surface and subsequent electrochemical characterization of the auto-inserted membrane protein. Therefore, we expect LAiR to be broadly utilized as a new platform for efficient investigation of IMPs in a lipid environment.

## Results

### Auto-insertion into liposomes

Traditionally, two methods have seen popularity for the removal of detergents to reintegrate detergent-solubilized IMPs into liposomes. Either biobeads are added successively or a rapid dilution (typically 200 fold) are used^7,8^. Both methods effectively leach detergent from the reintegration reaction to minimize detergent in the resulting proteoliposome preparation. The most sensitive proteins to detergent are multi-subunit IMPs in which lipid plays an important stabilizing role^3^. Even more sensitive and difficult to handle are IMPs that interact with hydrophobic substrates such as quinones^17,18^. While investigating the quinone-binding protein cytochrome *bo*_3_ (c*bo*_3_) from *Escherichia coli*, we observed that this IMP readily inserted into liposomes without biobead addition (Fig. 1a). c*bo*_3_ solubilized in dodecyl-maltoside (DDM), currently the most widely used detergent for IMP investigation^16^, auto-inserted into unilamellar liposomes (400 nm diameter) composed of *E. coli* polar lipids with >90 % efficiency after 30 min incubation (Fig. 1b). For comparison, reintegration using the standard biobead method or the rapid dilution approach yielded an efficiency of ~75 % or ~65 % respectively (Fig. 1b). Intriguingly, the time course of protein insertion into the liposomes revealed that c*bo*_3_ auto-inserted almost completely within <5 min (Fig. 1c), whereas the total time needed for reintegration with biobeads or using the rapid dilution method exceeded one hour.

**Fig. 1.**
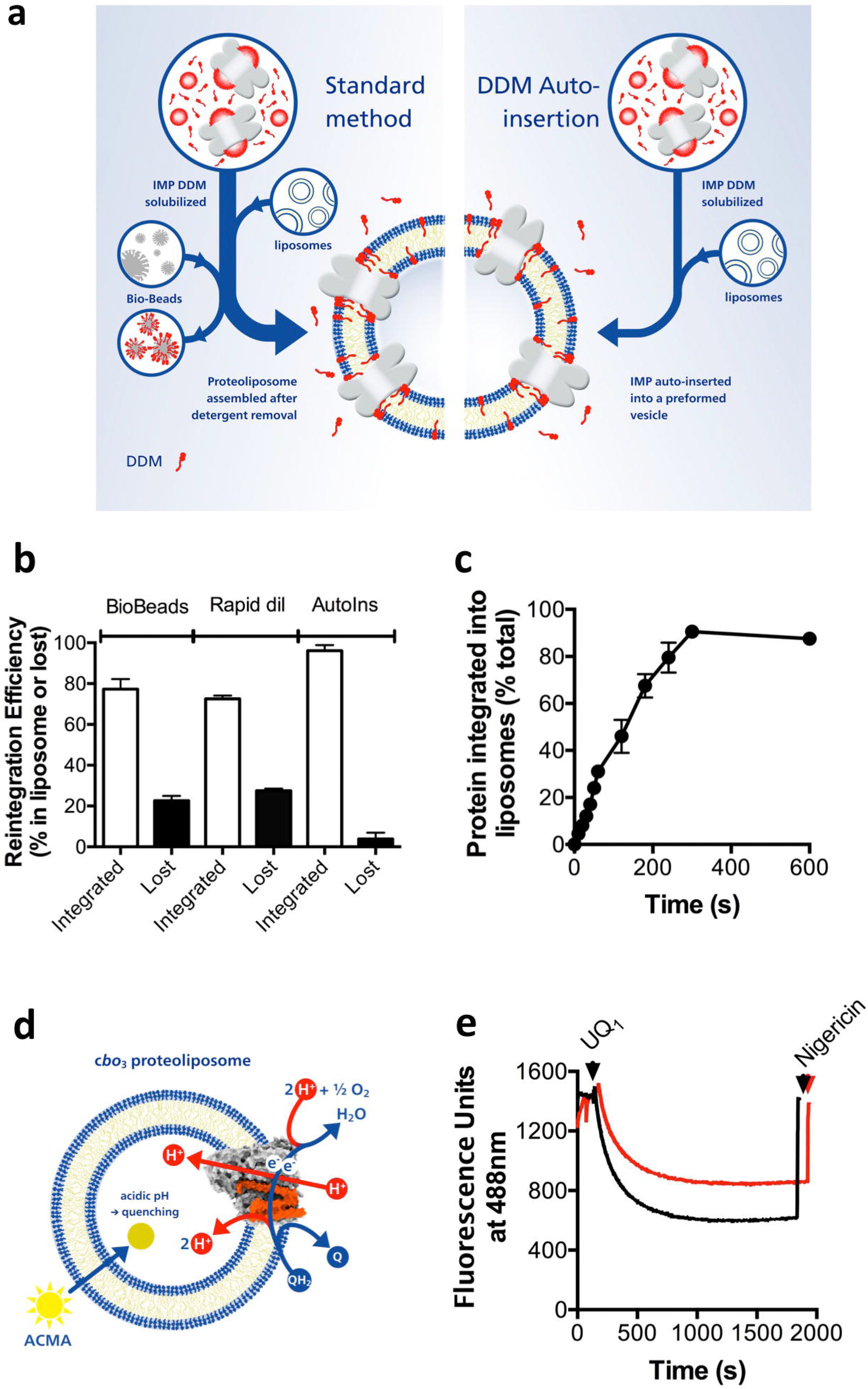
Auto-insertion is a rapid and highly efficient reintegration methodology. **a**, Schematic view of the standard reintegration method, where biobeads are used to reduce detergent concentration, in comparison with the DDM auto-insertion experimental system, where detergent is not removed. **b**, Reintegration efficiency (100% represents the total protein amount present in the sample) for DDM-purified c*bo*_3_ into *E. coli* polar lipids using either biobeads (>1.5 hrs), rapid dilution (>1 hr), or auto-insertion (30 min). **c**, Time course of reintegrating c*bo*_3_ into *E. coli* polar lipids using auto-insertion. **d**, Scheme of the experimental system used in panel **e** to assess proton pumping activity by reintegration c*bo*_3_. The reaction was started by addition of soluble ubiquinone-1 (UQ1) that is reduced by addition of dithiothreitol. As protons are pumped into the proteoliposome lumen, the membrane-permeable fluorescent dye ACMA is accumulated inside the vesicles, which causes quenching of its fluorescence. **e**, Time course of ACMA quenching. c*bo*_3_ proteoliposomes were obtained using either the standard biobeads method (black curve) or using DDM auto-insertion (red curve). Nigericin is an H^+^/K^+^ antiporter that offsets any proton concentration differences across the proteoliposome membrane. For each experiment 3 biological replicates were used, shown are either average values with standard deviation (**b, c**) or representative results (**e**).

Next, we assessed the enzymatic activity of the reintegrated protein. c*bo*_3_ is an enzyme that transfers electrons from a quinol-type substrate onto molecular oxygen to form water. This membrane-based electron transfer activity is coupled to trans-membrane proton pumping^19^ (Fig. 1d). We monitored the acidification of the proteoliposome lumen using a pH-sensitive fluorophore (Fig. 1d). Auto-inserted c*bo*_3_ clearly displayed proton pump activity, however, the degree of proteoliposome acidification was lower as compared to the biobead-reintegrated sample (Fig. 1e). These results show that auto-insertion can be utilized as a rapid, simple and convenient technique for reintegration of a DDM-solubilized IMP, but clearly the addition of biobeads reduces the extent of proton leak, leading to the hypothesis detergent may be a critical factor.

### <u>L</u>MNG <u>A</u>uto-<u>i</u>nsertion <u>r</u>eintegration (LAiR) provides optimal proton-tightness of proteoliposomes

The detergent, lauryl maltose neopentyl glycol (LMNG), has been described as suitable for preserving IMP stability and activity^20^ and for preparing IMPs for structural analysis by X-ray crystallography^20^ and cryo-electron microscopy^21^. LMNG’s lipid like molecular architecture conveys it with an extraordinarily low critical micelle concentration^20^ and high affinity for IMPs in solution^22^. These properties allow as little as 0.002 % LMNG to retain an IMP in solution (compared to ~0.025 % for DDM), a clear advantage if, as we hypothesized, proton leakage is resulting from residual detergent in the reconstituted proteoliposomes. Therefore, we attempted **L**MNG **A**uto-**i**nsertion **r**eintegration (**LAiR**; Fig. 2a). c*bo*_3_ was purified with LMNG and then auto-inserted into *E. coli* polar lipid liposomes with high rate and efficiency as observed for the DDM-reintegrated enzyme (Fig. 2b).

**Fig. 2.**
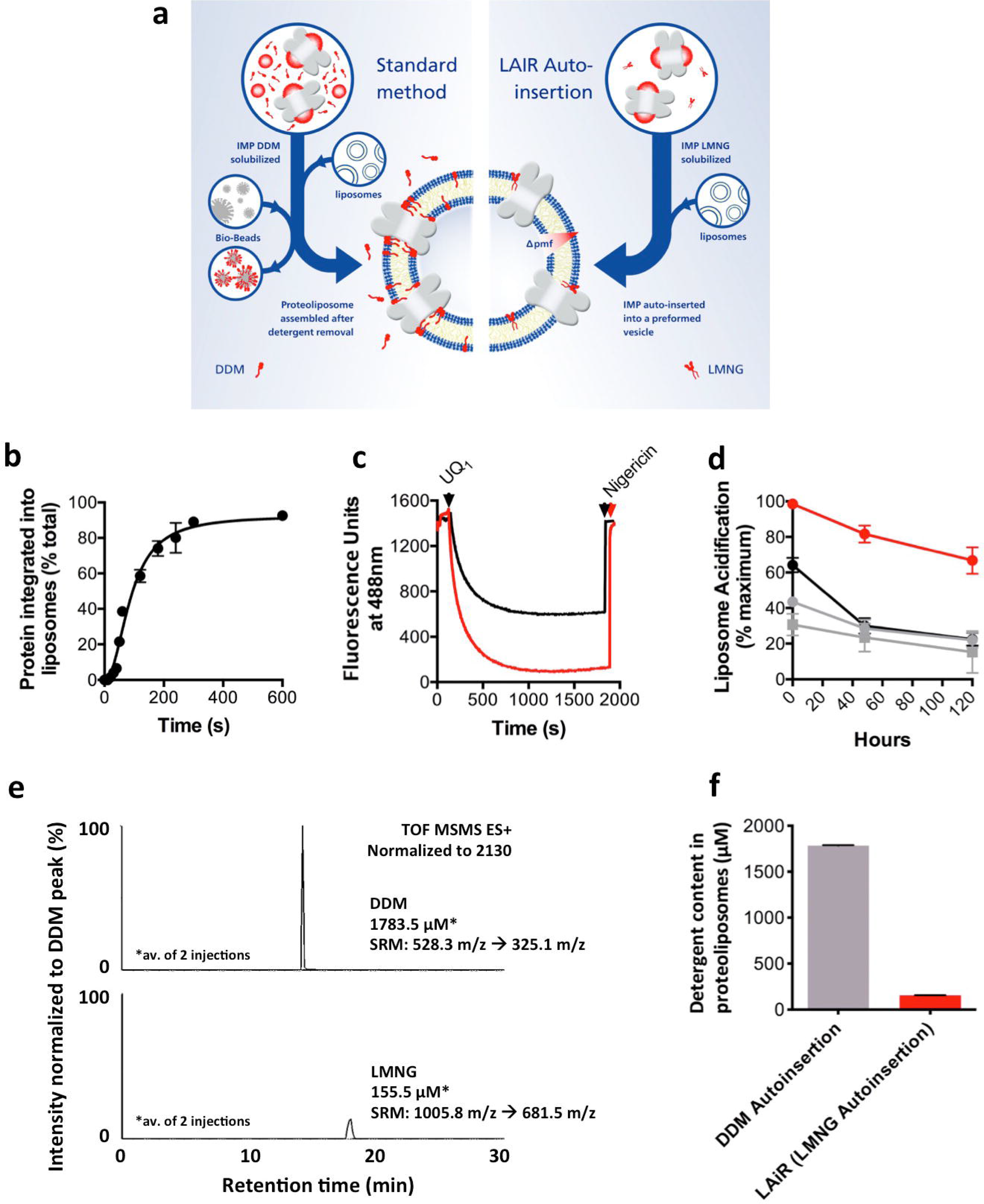
LMNG Auto-insertion Reintegration (LAiR) results in highly stable, proton-tight proteoliposomes. **a**, Schematic view of the standard reintegration method, where biobeads are used to reduce detergent concentration, in comparison with the LAiR method, where the ultra-low CMC detergent LMNG is used. **b**, Time course of reintegrating c*bo*_3_ purified using LMNG into *E. coli* polar lipids liposomes using auto-insertion. **c**, Proton-pumping activity of reintegrated c*bo*_3_ monitored by quenching of the ACMA fluorescence. Ubiquinol-1 (UQ_1_H_2_) was used to start the reaction. c*bo*_3_ proteoliposomes were prepared using either biobeads (black curve) or using LAiR after a 30 min time period (red curve). **d**, Cytochrome *bo*_3_ proteoliposome lifetime. Proteoliposomes were produced by either LAR (red line, circles), DDM auto-insertion (grey line, circles), by the biobeads/DDM method (black line, circles), or by rapid dilution reconstitution (grey line, squares). The proteoliposomes were stored on ice at 4°C over a period of 120 h (5 days). ACMA quenching assays were performed on days 1, 3 and 5 to assess proteoliposome quality versus time. The maximum degree of quenching was calculated and presented relative to the LMNG auto-insertion sample on day 0. **e** and **f**, Quantification of detergent remaining after completion of either DDM auto-insertion or LAiR. **e**, Residual detergent in proteoliposomes was quantified using LC/MS (selected reaction monitoring). The upper panel shows residual detergent from DDM auto-insertion and the lower panel shows residual detergent from LAiR. **f**, is a graphical comparison. For details, please refer to Online Methods. For each experiment 3 biological replicates (**a**, **c**), 2 technical replicates (**e**, **f**), or representative results (**b, d**) are shown with average value’s with error as a standard deviation of the mean.

Next, we explored the impact of the type of detergent empolyed on the proton tightness of the reconstituted proteoliposome. c*bo*_3_ proteoliposomes prepared using LAiR were capable of maintaining a 40 % higher level of acidification as compared to the biobead-inserted standard (Figure 2c). Proteoliposome acidification caused by pumping of protons into the lumen was completely reverted by addition of the H^+^/K^+^ antiporter nigericin (Fig. 2c), in line with a highly proton-tight proteoliposome system. Notably, c*bo*_3_ proteoliposomes prepared using LAiR also displayed high stability over extended times, with ~70 % of the maximal level of acidification maintained after incubation on ice for five days (Fig. 2d). In contrast, five-day incubation of c*bo*_3_, reintegrated by either DDM auto-insertion, biobeads, or rapid-dilution resulted in only ~20 % of the maximum activity measured for c*bo*_3_ proteoliposomes prepared using LAiR (Fig. 2d).

To test our hypothesis that it is the difference in the amount of residual detergent in the proteoliposomes that is underlying the significant higher proton tightness, we used LC/MS to quantify both the amount of residual DDM and LMNG after either DDM auto-insertion or LAiR. For the biobead/DDM sample, residual biobeads precluded an accurate detergent quantification. Indeed, we found over 10-fold lower amount of residual detergent for c*bo*_3_ proteoliposomes prepared using LAiR as compared to the proteoliposomes prepared using DDM auto-insertion (Figs. 2e and 2f).

### LAiR retains high membrane-bound activity under control of the proton motive force

Next, we evaluated the ability of LAiR to investigate IMP properties that are currently out of reach of standard reintegration methods. As an example, we examined the impact of the proton motive force across the membrane on the electron transfer activity of c*bo*_3_. The proton motive force is a key effector of c*bo*_3_ membrane-bound activity, however, insufficient proton tightness of proteoliposomes precluded investigation in classical systems^23^.

We used LAiR to reintegrate *E. coli* c*bo_3_* into *E. coli* polar lipid liposomes and immobilized the resulting proteoliposomes on a 6-mercaptohexanol (6MH) modified gold electrode surface *via* a self-assembled monolayer (SAM, Fig. 3a). As a control, we reintegrated *E. coli* c*bo*_3_ using the biobead standard method. To assess the membrane-bound electron transfer activity by c*bo_3_* we used cyclic voltammetry (Fig. 3a). We found that c*bo_3_* proteoliposomes prepared using LAiR displayed approximately 3-fold higher maximal activity than the control reintegrated using the biobead standard method (Fig. 3b,c). Proteoliposomes prepared using LAiR thus display not only enhanced lumen acidification (as shown in Fig. 2b), but the reintegrated enzyme also retains higher membrane-bound activity as compared to c*bo*_3_ reintegrated using the biobead standard method.

**Fig. 3.**
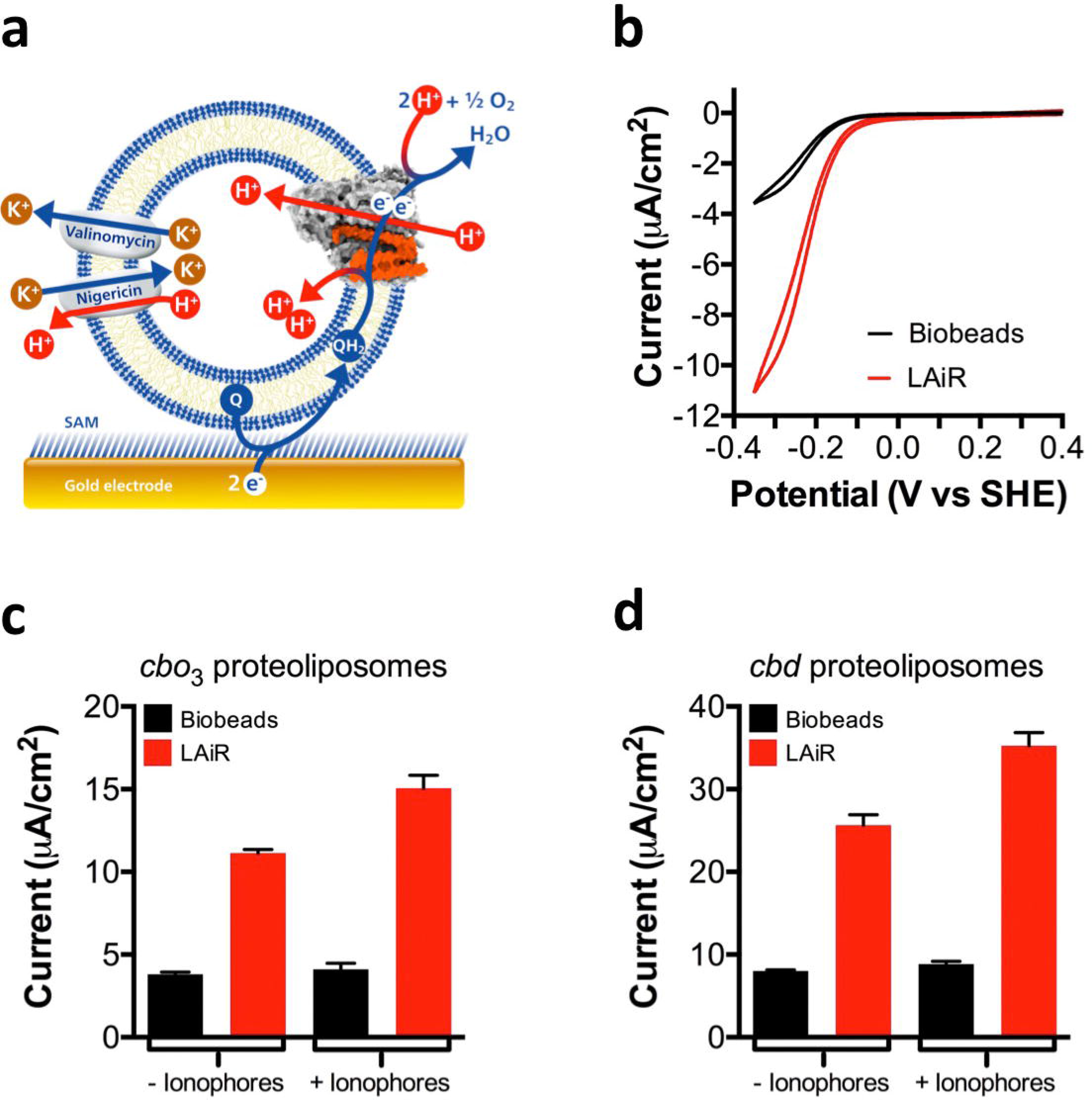
LAiR visualizes control of membrane-based IMP activity by the proton motive force. **a**, Scheme showing the experimental set-up used to study membrane-based electron transfer by LAiR-inserted c*bo*_3_. Proteolipomes containing c*bo*_3_ are immobilized *via* a 6-mercaptohexanol self-assembled monolayer (SAM) onto an electrode. The electrode feeds in electrons that reduce a membrane-bound quinone to quinol at the reduction potential of ubiquinone-10 (UQ_10_). The proton motive force generated by c*bo*_3_ can be dissipated by two ionophores, the potassium channel valinomycin together with the H^+^/K^+^ antiporter nigericin. **b**, Cyclic voltammetry with the immobilized c*bo*_3_ proteoliposomes, showing a comparison of electron transfer activity between LAiR (red curve) and biobead/DDM based reintegration (black curve). **c**, Impact of the proton motive force on electrochemically detected oxygen consumption activity by cytochrome *bo*_3_ (c*bo*_3_) proteoliposomes. **d**, Impact of the proton motive force on electrochemically detected oxygen consumption activity by cytochrome *bd* (c*bd)* proteoliposomes. For each experiment 3 biological replicates were used. Shown are either representative results (**b**) or average values with standard deviation (**c,d**).

We then evaluated the effect of the proton motive force on the electron transfer activity, a property that has not been measurable to date in the same system^23^. To dissipate the proton motive force, we added a combination of two ionophores: the K^+^ channel valinomycin and the H^+^/K^+^-antiporter nigericin. The electron transfer activity of the biobead reintegrated c*bo_3_* did not noticeably change upon addition of valinomycin/nigericin (Fig. 3c), indicating that in these proteoliposomes c*bo*_3_*-* catalyzed electron transfer is not highly regulated by the proton motive force within the time window measured. In contrast, the electron transfer activity of the c*bo_3_* proteoliposomes prepared using LAiR considerably increased after the addition of valinomycin/nigericin, demonstrating that the membrane-based electron transfer activity of c*bo_3_* is under tight control of the proton motive force (Fig. 3c).

To exclude that control of membrane-bound activity by the proton motive force represents an idiosyncratic property of c*bo*_3_ proteoliposomes prepared using LAiR, we extended our efforts to cytochrome *bd* (c*bd*) from *E. coli*. c*bd* is a quinol oxidase that reduces oxygen to water and contributes to the proton motive force by vectorial uptake and release of protons, but it is not evolutionary related to c*bo*_3_^24^. Similar to the results obtained for c*bo*_3_, the membrane-bound electron transfer activity of the biobead reintegrated c*bd* displayed only minimal change upon dissipation of the proton motive force within the time window measured, whereas the activity of c*bd* proteoliposomes prepared using LAiR considerably increased, revealing tight control by the proton motive force (Fig. 3d). Taken together, the results demonstrate that auto-insertion allows for investigation of IMPs in the context of an ion motive force, mimicking the native non-equilibrium biomembrane environment in a living cell.

### LAiR allows non-destructive reintegration of an IMP into a pre-existing planar membrane

Solid-supported lipid membranes and planar bilayers allow for the study of IMPs by analytical tools such as atomic force microscopy, impedance spectroscopy, surface plasmon resonance sensing, and quartz-crystal microbalance^25^. Currently, to investigate IMPs in these surface-tethered bilayers, the proteins are first reintegrated into liposomes and in a second step the resulting proteoliposomes are disrupted to form a bilayer on a gold electrode surface modified with a tethering SAM (Fig. 4a)^26^. In order to further evaluate the scope of the auto-insertion approach, we explored its applicability to pre-existing membrane bilayers tethered onto an electrode surface (Fig. 4b). To demonstrate this utility, we assembled a tethered membrane bilayer using liposomes composed of *E. coli* polar lipids containing substrate and then used LAiR to reintegrate c*bo*_3_ (Fig 4b). As a control, we reintegrated c*bo*_3_ into liposomes of identical composition using a standard biobead protocol and then assembled a tethered membrane bilayer using these proteoliposomes (Fig. 4a).

**Fig. 4.**
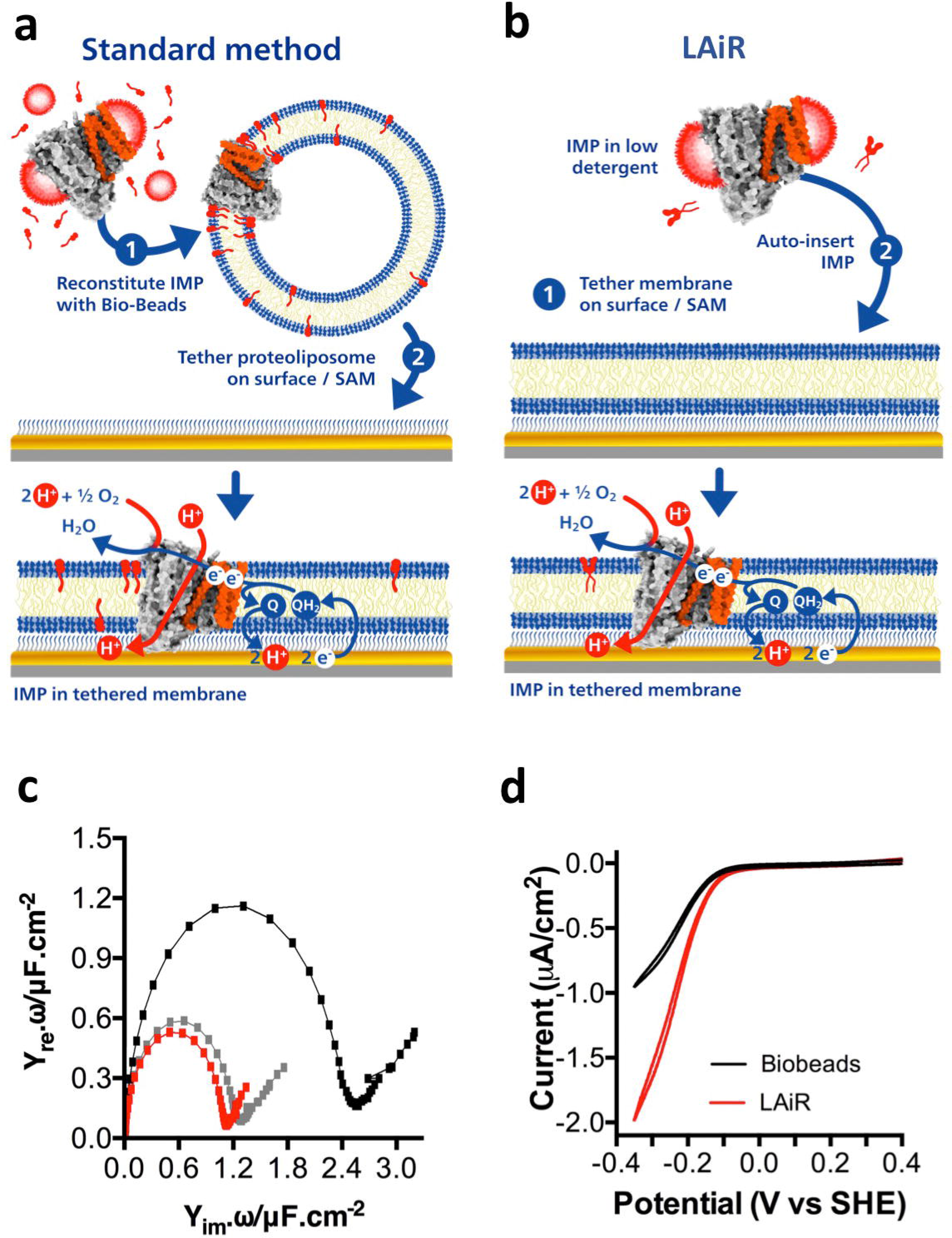
Using LAiR for integration of IMPs into pre-existing planar membranes. **a** and **b**, Schemes for integrating an IMP into a surface-tethered membrane (tBLM) using a standard method or LAiR. To assess enzymatic activity, the electrode feeds in electrons that reduce a membrane-bound quinone to quinol at the reduction potential of ubiquinone-10 (UQ_10_; displayed as ‘Q’). **a**, Using the standard method, an IMP is first reintegrated into a liposome containing UQ_10_ using either biobeads or rapid dilution, and the resulting proteoliposome is chemically disrupted onto a specialized tethering self-assembled monolayer (SAM). **b**, Using LAiR, a membrane bilayer is first tethered onto the surface *via* the self-assembled monolayer in an identical method to **a**, but without reintegrated protein. Subsequently, the IMP is auto-inserted into the tethered membrane bilayer. **c**, Impedance spectroscopy of the system shown in **b**. The black curves show the impedance of the SAM before the membrane bilayer was added, the grey curve shows impedance after membrane bilayer formation, and the red curve is impedance after LAiR of c*bo*_3_. The size of the half circle is indicative of the accessibility of solution phase ions to the electrode. A large half circle indicates easy access to the electrode (i.e. no membrane or holes in a membrane) while a small half circle indicates restricted access (i.e. a tight membrane). **d**, Cyclic voltammetry comparing between the electrochemically detected oxygen consumption activity of a c*bo*_3_ in a tBLM formed by the biobead method (black curve) or by LAiR (red curve). For each experiment 3 biological replicates were used, representative results are shown.

To evaluate membrane protein reintegration, we assessed the permeability of the tethered membrane using impedance spectroscopy. Formation of the tethered membrane bilayer decreased the permeability as revealed by a drop in capacitance (Fig. 4c). Interestingly, upon reintegration of c*bo*_3_ using LAiR the capacitance marginally decreased further (Fig. 4c), suggesting that protein integration into the pre-formed membrane was not only non-destructive, but that the protein appeared to tighten the membrane. Cyclic voltammetry revealed that the activity of *cbo*_3_ reintegrated using LAiR into the pre-formed tethered membrane was two-fold higher than that observed with a membrane formed using a membrane assembled from c*bo*_3_ proteoliposomes prepared by biobead reintegration (Fig. 4d), confirming highly effective c*bo*_3_ insertion. This demonstrates that LAiR not only can integrate protein into a pre-formed tethered membrane in a rapid, non-destructive manner, but also that the auto-inserted protein’s activity is significantly higher than that in current state-of-the-art experimental set-ups. As such, LAiR can serve as a versatile method to integrate IMPs into surface-based biosensors.

### LAiR preserves intactness and higher-order oligomeric features of a fragile multi-subunit IMP

To evaluate if LAiR can also be utilized for fragile and highly complex multi-subunit IMPs of mammalian origin, we assessed the impact of LAiR on the intactness of F-type ATP synthase from bovine heart mitochondria. The mammalian F-type ATP synthase is a large (MW >600 kDa) IMP complex comprising 28 subunits in its monomeric form. It consists of two distinct domains, F_1_, which catalyzes ATP hydrolysis (or synthesis, depending on the direction of the reaction), and F_o_, which, coupled to ATP hydrolysis, conducts protons across the membrane (Fig. 5a)^27^. In mammalian mitochondria, the F-type ATP synthase forms oligomeric supercomplexes of rows of dimers which are located at the ridge of cristae and bend the membrane by ~85 degree, i.e. represent a crucial molecular player in cristae architecture^28,29^. We purified LMNG-solubilized F-type ATP synthase from bovine heart mitochondria^30^ and used LAiR to reintegrate it into liposomes prepared from bovine heart lipids. Similar to c*bo*_3_, preparation of F-type ATP synthase proteoliposomes using LAiR was virtually completed within several minutes (Fig. 5b). Negative stain EM and cryo electron tomography revealed the F-type ATP synthase complexes to be densely packed in the liposomes and indicated their structural integrity (Fig. 5c and 5d, Sup. Movies 1 and 2). The resulting proteoliposomes maintained high proton tightness over several days (Fig. 5e). These results demonstrate that LAiR is not restricted to prokaryotic systems but is also applicable to IMPs and lipids extracted from eukaryotic sources.

**Fig. 5.**
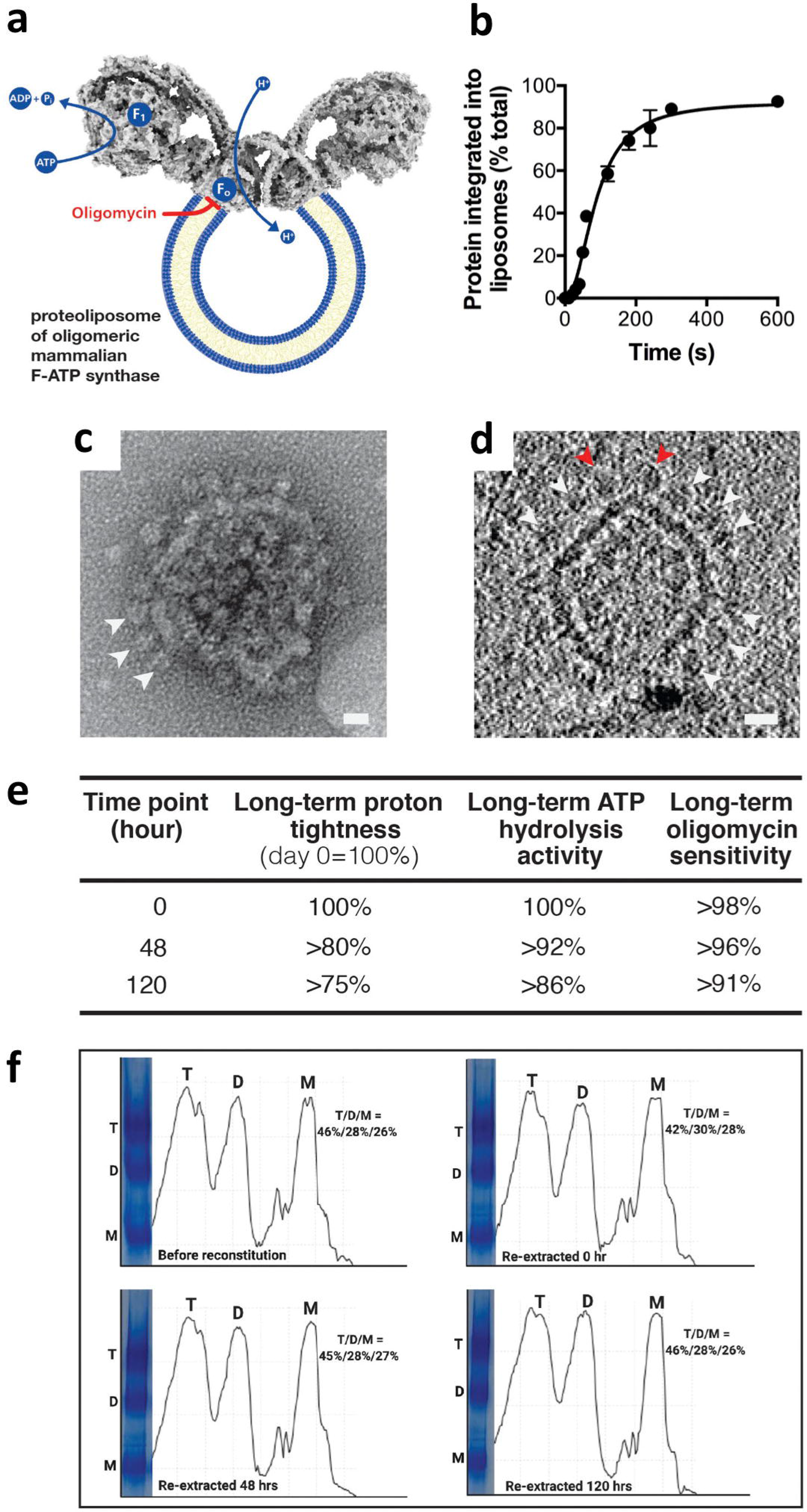
Structural integrity and inhibitor-sensitivity of a multi-subunit IMP is maintained by LAiR. **a**, Scheme of a mammalian F-type ATP synthase dimer and the reactions it catalyzes. The F_1_ part, where ATP hydrolysis occurs, and the F_o_ domain, where the inhibitor oligomycin binds, are indicated. **b**, Time-course of reintegration. F-type ATP synthase purified from bovine heart mitochondria was reintegrated into liposomes composed of bovine heart lipids. **c** and **d**, Reintegration of F-type ATP synthase complexes into the proteoliposomes as assessed by negative-stain (**c**) or cryo-electron tomography after a 30 min LAiR treatment. **c**, Shows a negative-stained intact proteoliposome, and **d**, shows a cryo-tomograph section of a proteoliposome (also see Sup. Movie 1). White arrows indicate F-type ATP synthase complexes decorating the proteoliposomes in both **c** and **d**, while red arrows in **d** indicate the presence of an intact membrane bending F-type ATP synthase oligomer. **e**, Long-term proteoliposome tightness and functional coupling between the F_o_ and F_1_ domains of the reintegrated F-type ATP synthase. The proteoliposomes with reintegrated ATP synthase were stored on ice at 4°C over a period of 120 h (5 days). The proton tightness was evaluated by measuring acidification of the liposome lumen upon hydrolysis of ATP using the fluorescent dye ACMA. Quenching of ACMA fluorescence is presented for each time point relative to quenching at time point 0. The functional coupling between the F_o_ and F_1_ domains was measured by the F_o_ inhibitor oligomycin A. ATP hydrolysis was measured coupled to NADH oxidation (via the pyruvate kinase and lactate dehydrogenase reactions), whose absorption at 340 nm was monitored. **e**, Long-term stability of reintegrated F-type ATP synthase dimers and tetramers. After LAiR, the F-type ATP synthase was re-extracted from the proteoliposomes at the indicated time points and subjected to Native PAGE. For each experiment, at least 2 biological replicates were used. Average values with standard deviation (**b, e**) or representative results (**c, d, f**) are shown.

A key problem in the investigation of F-type ATP synthase is that upon exposure to detergents, during liposome reintegration or upon long-term storage, the enzyme is prone to a functional disconnection between its two domains, thereby uncoupling proton flow in the F_o_ domain from ATP hydrolysis in the F_1_ domain^27^. As a consequence, inhibitors of F_o_ proton transport typically only partially prevent ATP hydrolysis by the F_1_ domain^31^. We measured ATP hydrolysis activity using an assay that connects hydrolysis of ATP to oxidation of NADH, allowing for optical read-out. We found that the ATP hydrolysis activity by the auto-inserted F-type ATP synthase was almost completely (>98 %) blocked by oligomycin A, an inhibitor of the F_o_ domain (Fig. 5a and 5e). The high degree of oligomycin sensitivity was maintained even after 5-day incubation on ice (Fig. 5e). These results reveal strong functional coupling between the F_1_ and F_o_ domains and demonstrate that a fragile multi-subunit IMP can be stably reintegrated into liposomes using LAiR and remain intact over extended periods. F-type ATP synthase inhibitors currently are under development for various applications, e.g. as anti-bacterials^32^, against age-associated dementia^33^, as anti-cancer agents^34^ and ischemia/reperfusion^35^ The functionally intact, stably LAiR-reintegrated F-type ATP synthase, combined with optical read-out, has a high potential as a valuable test-system for these efforts. These results indicate that auto-inserted multi-subunit IMPs can be utilized in inhibitor screening systems, where accurate, quantitative characterization of prospective drugs is imperative and long-term usage can be required.

A second key feature of mitochondrial F-type ATP synthase, itself a multi-subunit complex, is its ability to form higher-order assemblies such as dimers or tetramers in the inner mitochondrial membrane^28^. Higher-order structural features can strongly influence the functionality of an IMP. In case of mitochondrial F-type ATP synthase formation of dimers is instrumental for folding the cristae structure of the inner membrane^28,29^, a key factor in mitochondrial physiological function. However, mitochondrial F-type ATP synthase is prone to dissociation into monomers in the presence of DDM^36^ and proper reconstitution of purified dimeric F-type ATP synthase is challenging^37,38^. We confirmed that LAiR reintegrated of bovine heart F-type ATP synthase did not cause noticeable dissociation of dimers or tetramers into monomers (Fig. 5d and 5f). The monomer/dimer/tetramer ratio of the auto-inserted enzyme was actually maintained over a period of 5 days when stored on ice at 4°C (Fig. 5f). As such, auto-insertion may facilitate long-term investigation of complex mammalian IMPs without altering their higher-order oligomeric state.

## Conclusion & Outlook

Our results demonstrate that LAiR reintegration of IMPs maintains even fragile IMPs in an active, stable state, and allows detection and study of IMP properties that are not accessible with standard methods. The functional intactness exhibited by proteoliposomes comprising IMPs and lipids from prokaryotic or eukaryotic sources opens hitherto closed doors to the investigation of a broad scope of problems in biomembrane research. The long-term ion-tightness associated with LAiR enables the researcher to perform work at ease, spreading experimentation on an individual sample over several days with intermediate storage on ice. As the approach is technically straightforward, LAiR can likely be adopted in various laboratories without extensive method-specific training.

The molecular processes during LAiR and the factors responsible for the observed proteoliposome proton-tightness and high enzymatic activity are not well understood yet. However, it is plausible that the almost complete absence of free detergent micelles in solutions of IMPs stabilized by 0.002% LMNG together with the lipid-like molecular architecture of LMNG are key factors that enable the extraordinary tightness and stability of LAiR proteoliposomes^21,39^. A systematic comparison of different detergents or detergent-derivatives, e.g. using derivatives that have been reported for LMNG^20^ or can be synthesized in the future, may provide clues on the underlying molecular mechanism, and may further improve the performance of this new method.

LAiR will broaden the utility of membrane biochemistry and may also prove useful in fields such as electrochemistry, electrophysiology and cryo-EM of proteoliposomes^38^. Next to the importance for fundamental research, the high stability of proteoliposomes comprising LAiR reintegrated multi-subunit IMPs can also enable long-term usage in applied settings, e.g. in drug discovery screens or in biosensor systems based on surface-tethered membranes. Finally, LAiR also has a strong potential to provide input for efforts in synthetic biology aimed at constructing artificial, liposome-based systems mimicking cell-like functions^40^, and might even allow for integration of IMPs into native cell membranes to modify cell function.

## Acknowledgements

We would like to thank Hiroyuki Noji for support for the initial stages of this project, and both Hiroyuki Noji and Naoki Soga for insightful discussions. A.H.A. wishes to thank the Royal Embassy of Saudi Arabia (NL) and King Abdulaziz University in Saudi Arabia for financial support.

## Funding

This work was supported by a long-term International fellowship from the Japan Society for the Promotion of Sciences (JSPS; P14383) to D.G.G.M; Delft University of Technology Start-up Grant to D.G.G.M; a BINDS grant from AMED (JP16K07266) to Atsunori Oshima and C.G; a Grants-in-Aid for Scientific Research (B) (JP 17H03647) from MEXT to C.G; the International Joint Research Promotion Program from Osaka University to Genji Kurisu and C.G; a JST-CREST Grant Number (JP18071859) to K.M; the Naito Foundation Subsidy for Female Researchers after Maternity Leave (C.J) and JSPS 25-5370 (C.J).

## Author Contributions

A.H.A., A.G-H, A.P., C.J., C.d.R., M.P., C.G., K.M., D.G.G.M performed experiments; H.L. provided expertise and data; M.P, C.G., D.B., and D.G.G.M supervised experiments; C.G., D.B., and D.G.G.M. supervised the over-all research and wrote the manuscript with contributions from all co-authors.

## Competing Interests statement

The authors declare no competing financial interests.

## Online Methods

### Chemicals

β-dodecyl-maltoside (DDM, Anagrade, < 0.2 % α-configuration) and Lauryl maltose neopentyl glycol (LMNG) were purchased from Anatrace. Lipids were purchased from Avanti Polar Lipids. Other chemicals were obtained from Sigma.

### Purification of cytochrome *bo_3_* from *Escherichia coli*

Cytochrome *bo*_3_ (c*bo*_3_) was extracted and purified from inner membranes based on Hards et al. 2018^1^. *E. coli* was aerobically grown to mid-log phase at 37°C in Luria-Bertani (LB) medium supplemented with 500 μM CuSO_4_ and 100 μg ml^−1^ carbenicillin. Cells were harvested by centrifugation at 10,000 × *g* for 10 mins. The pellet was then washed, and repelleted twice with buffer B (20 mM 3-N-morpholinopropanesulfonic acid (MOPS), 30 mM Na_2_SO_4_, 10% Glycerol pH 7.4). Cells were then resuspended in buffer B containing 1 Roche cOmplete™ mini protease inhibitor tablet per 50 mL, 0.1 mM phenylmethylsulfonyl fluoride, 0.1 mg pancreatic DNase per mL, and lysed by two passages through a French press at 20,000 psi. Any remaining debris and unbroken cells were removed by centrifugation at 10,000 × *g* for 30 min. The supernatant was then ultracentrifuged (200,000 × *g*, 45 min, 4°C) and the membrane pellet resuspended in buffer C (20 mM MOPS, 30 mM Na_2_SO_4_, 25% w/w sucrose, pH 7.4). This was applied to the top of a 30% w/w to 55% w/w sucrose gradient and ultracentrifugation (130,000 × *g*, 16 h, 4°C) with no deceleration or breaking to separate inner membrane from outer membrane. The inner membrane fraction was removed from the sucrose gradient and washed 3 times with buffer B by ultracentrifugation (200,000 × *g*, 45 min, 4°C). Inner membranes were then resuspended in buffer B and either used immediately for purification or stored in aliquots at −80°C until use. To extract c*bo*_3_, inner membrane vesicles were diluted to 5 mg of protein mL in solubilization buffer (20 mM Tris HCl, pH 8.0, 5 mM MgSO_4_, 10% glycerol, 0.5% LMNG or 1% DDM, 300 mM NaCl, 10 mM imidazole) and incubated at 30°C for 30 min with gentle inversion every 5 min. The unsolubilized material was removed by ultracentrifugation (180,000 × *g*, 45 min, 4°C), and the supernatant was applied to a Nickel-Sepharose High Performance (GE Healthcare) column that was previously washed with water and equilibrated with IMAC buffer (50 mM Tris HCl, pH 8.0, 5 mM MgSO_4_, 10% glycerol, 0.005% LMNG or 0.02% DDM, 300 mM NaCl) containing 10 mM imidazole. To remove contaminating proteins, the resin was washed with IMAC buffer containing 30 mM imidazole and 150 mM NaCl, and c*bo*_3_ was eluted with IMAC buffer containing with 200 mM imidazole, 150 mM NaCl, and 20% glycerol. The red c*bo*_3_ containing fractions were pooled and concentrated using an Amicon Ultra centrifugal filter device (molecular weight cutoff ((MWCO), 100,000).

### Purification of cytochrome *bd* from *Escherichia coli*

Cytochrome *bd* (c*bd*) was purified from *E. coli* based on Goojani *et al*.^2^. Briefly, *E. coli* strain MB43 carrying the pET17cydABX-Strep-tag plasmid was grown in Luria-Bertani medium with 100 μg/ml Ampicillin at 37°C overnight with shaking at 200 rpm. The bacteria were diluted to OD_600_ 0.01 in 800 ml LB medium with 100 μg/ml Ampicillin and incubated until reaching OD_600_ 0.4. Then IPTG (0.45 mM final conc.) was added and the bacteria were incubated again at 37°C, 200 rpm until reaching OD_600_ 2.0. Cells were sedimented by centrifugation at 6,000 × *g* for 20 min (JA-10 rotor). The pellets were washed by phosphate buffer saline, pH 7.4, and pelleted at 6,000 × *g* for 20 min. Approximately 15 g of wet cells were re-suspended with 75 ml of MOPS solution (50 mM MOPS, 100 mM NaCl and protease inhibitor (C0mplete EDTA free)). The cells were disrupted by three passages though a Stansted cell homogenizer at 18,000 psi. Unbroken cells were centrifuged at 9,500 × *g* for 20 min. Subsequently, the supernatant was pelleted by ultracentrifugation 250,000 × *g* for 75 min at 4°C. The pellet was re-suspended in MOPS solution and the protein concentration was determined To extract c*bd*, the protein concentration was adjusted to 10 mg/ml with MOPS solution and either DDM (1% final conc.) or LMNG (0.5% final conc.) were added and the solution was incubated at 4°C for an hour with gentle shaking. Un-solubilized material was sedimented by ultracentrifugation at 250,000 × *g* at 4°C for 15 min (70-Ti rotor). The collected supernatant was applied on streptactin column at 4°C. To remove nonspecifically bound protein, the column was washed with 50 mM sodium phosphate pH 8.0 containing 300 mM NaCl, protease inhibitor (cOmplete EDTA free), and 0.01% DDM or 0.005% LMNG. Finally, the protein was eluted from the column with 50 mM sodium phosphate pH 8.0 containing 300 mM NaCl, protease inhibitor (cOmplete EDTA free), 0.01% DDM or 0.005% LMNG, and 2.5 mM desthiobiotin.

### Purification of F-type ATP synthase from *Bos taurus* (Bovine)

Purification of mammalian F-type ATP synthase was conducted as previously described^3,4^. Fresh *B. taurus* hearts were obtained immediately after slaughter by an authorized slaughterhouse and inner mitochondrial membranes were purified according to Shimada et al.^5^. Fat and connective tissues were carefully removed allowing the preparation of 1,000 g of minced meat. Each 500 g was suspended in 3,250 ml of 23 mM sodium phosphate buffer (pH 7.4 at 0°C) and homogenized for 5 min at 13,000 rpm in a homogenizer (Nihon Seiki), followed by centrifugation for 20 min at 2,800 rpm in a refrigerated centrifuge (Kubota Model 9810; RS-6600 rotor). The precipitate was suspended in 3,375 ml of 22.2 mM sodium phosphate buffer (pH 7.4) and re-homogenized, followed by centrifugation for 20 min at 2,800 rpm as previously. Supernatants were then combined and centrifuged for a further 30 min at 10,000 rpm with a refrigerated centrifuge (Beckman Model Avanti HP-30I) using a JLA-10.500 rotor. The precipitate was then suspended in 50 mM Tris-HCl buffer (pH 8.0) and pelleted for 30 min at 30,000 rpm with an ultracentrifuge (Beckman Model-7) using a 45 Ti rotor. The pellet was suspended in 50 mM Tris-HCl buffer (pH 8.0) containing 660 mM sucrose to a final protein concentration of ~23 mg/ml. The suspension was kept in a 40 mM HEPES bufer (pH 7.8) containing 2 mM MgCl_2_, 0.1 mM EDTA, and 0.1 mM DTT and solubilized on ice *via* slow addition of deoxycholate and decylmaltoside to final concentrations of 0.7% (wt/vol) and 0.4% (wt/vol), respectively. The suspension was then centrifuged at 176,000 × *g* for 50 min and the supernatant applied to a sucrose step gradient (40 mM HEPES pH 7.8, 0.1 mM EDTA, 0.1 mM DTT, 0.2% wt/vol decylmaltoside and 2.0 M, 1.1 M, 1.0 M, or 0.9 M sucrose) and centrifuged at 176,000 × *g* for 15.5 hr. Fractions exhibiting ATP hydrolysis activity were loaded onto a Poros-20HQ ion-exchange column. The detergent was exchanged to LMNG using a double gradient from 0.2% to 0% decyl-maltoside and 0%–0.05% LMNG for 80 min at 1 ml/min. Complexes were eluted by a linear concentration gradient of 0–240 mM KCl in 40 mM HEPES pH 7.8, 150 mM sucrose, 2 mM MgCl_2_, 0.1 mM EDTA, 0.1 mM DTT and 0.005% (wt/vol) LMNG.

### Polyacrylamide gel electrophoresis (PAGE) and protein quantification

IMP preparations were routinely analyzed on 12% SDS-PAGE (data not shown, a detailed characterization of the LMNG-purified IMPs will be published elsewhere). Protein concentrations were determined using a bicinchoninic acid (BCA) protein assay kit (Sigma) with bovine serum albumin as the standard.

For Blue-Native page after LAiR, F-type ATP synthase was extracted from the proteoliposomes at the indicated time points using 0.5% LMNG. After centrifugation at 20,000 × *g* for 10 min at 4 °C, the supernatants were prepared for and separated by 3-12% Bis-Tris Mini-gels (Novex Life Technologies) according to the manufacturer’s instructions. ATP synthase bands were analyzed using ImageStudioLite software.

### Lipid treatment and proteoliposome preparation

Lipids used in this study were purchased from Avanti Polar Lipids, Inc., Alabaster, AL. Lipids were stored and treated as in McMillan *et al*^6^. Stock solutions of either native *E. coli* polar lipids extract (ECPL) or Bovine heart lipids (BHPL) were dried under an N_2_ stream. Where appropriate, ubiquinone (UQ_10_) was added to the chloroform-dissolved lipid mixtures at 1% mass/mass and dried together. Liposomes were resuspended in 20 mM Tris-(hydroxymethyl)aminomethane hydrochloride (Tris-HCl, Sigma) buffer containing 100 mM KCl. All liposome concentrations after resuspension were 10 mg/ml. These were then extruded 11 times through a 400 nm track edge membrane using an extruder (Avanti Polar Lipids, Inc., Alabaster, AL).

### Reintegration of IMPs into liposomes

#### Biobeads Method

Lipids (8 mg/ml) were solubilized in reintegration buffer (20 mM MOPS (pH 7.4) 30 mM Na_2_SO_4_, 100 mM KCl) containing 55 mM octyl glucoside by brief ultrasonic treatment. Purified IMP was added to a final concentration of 0.02 mg/ml, and the mixture was slowly stirred for 15 min at 20°C. Activated SM^2^ biobeads (80 mg wet beads/ml) were added directly to the mixture, and stirring continued for 30 min. After this, another 80 mg/ml biobeads was added followed by another 30 min incubation. Finally, another 160 mg/ml was added followed by a 90 min incubation to attempt to complete the removal of detergent. The top phase with the proteoliposomes was carefully removed with a pipette tip that did not pass any biobeads and diluted 10-fold with reintegration buffer and ultra-centrifuged at 180,000 × *g* for 60 min. The pellet resuspended in reintegration buffer, and used as indicated in figures.

#### Rapid dilution method

Lipids (8 mg/ml) were solubilized in reintegration buffer (20 mM MOPS (pH 7.4) 30 mM Na_2_SO_4_, 100 mM KCl) containing 55 mM octyl glucoside by brief ultrasonic treatment. Purified IMP was added to a final concentration of 0.02 mg/ml, and the mixture was slowly stirred for 60 min at 20°C. The mixture was then diluted 200-fold in reintegration buffer and ultra-centrifuged at 180,000 × *g* for 60 min. The pellet resuspended in reintegration buffer and used as indicated in figures.

#### DDM Auto-insertion method and LAiR

Lipids (10 mg/ml) were re-solubilized by vortex in reintegration buffer and extruded 13 times through a 400 μM polycarbonate membrane. Purified IMP was added to a final concentration of 0.02 mg/ml, and the mixture was slowly inverted for 5-30 min (as indicated in figures) at 20°C. The reconstitution was then either used immediately or diluted 10-fold and ultra-centrifuged at 180,000 × *g* for 30 min (as indicated). If the latter treatment was used, the pellet was resuspended in reintegration buffer and used as indicated in figures.

### Biochemical assays

#### ATP hydrolysis activity after reconstitution

ATP hydrolysis activity was measured at 38°C with stirring at 1,000 rpm using an ATP regenerating assay^7^. The assay mixture contained 50 mM MOPS (pH 7.4), 30 mM NaCl, 100 mM KCl, 3 mM phospho*enol*pyruvate, 1.5 mM MgCl_2_, 0.25 mM NADH, 1 μM valinomycin, 0.57 U/ml pyruvate kinase, 3.2 U/ml lactate dehydrogenase and 2 mM ATP. The reaction was initiated by the addition of 10 μg of auto-inserted ATP synthase into 1 ml of assay mixture. The rate of NADH oxidation was monitored continuously at 340 nm using a modified Cary 60 spectrophotometer (Agilent). Where indicated 2 μM oligomycin was added. The activity that hydrolyzed 1 μmol of ATP per min is defined as 1 unit. The activity of reconstituted ATP synthase at day 0 was 4.6 U/mg protein.

#### ATP-dependent liposome acidification (ACMA fluorescence quenching)

ATP-dependent proton translocation was determined at 38°C based on the quenching of ACMA. The 1.5-ml reaction mixture contained 50 mM MOPS (pH 7.4), 30 mM NaCl, 100 mM KCl, 3 mM phospho*enol*pyruvate, 1.5 mM MgCl_2_ 0.25 mM NADH, 0.57 U/ml pyruvate kinase, 3.2 U/ml lactate dehydrogenase, 1 μM ACMA, 1 μM valinomycin, and 10 μg F_1_F_o_ ATP synthase complexes reintegrated in bovine heart lipid liposomes. After the fluorescence signal had stabilized, the reaction was initiated by the addition of 2.5 mM neutralized ATP. Fluorescence was measured with an excitation wavelength of 410 nm and an emission wavelength of 480 nm (slit width, 10 nm) in a modified Cary Eclipse photospectrophotometer (Agilent). ATP-dependent quenching of ACMA fluorescence at day 0 was 75 %.

#### Quinone-dependent liposome acidification (ACMA fluorescence quenching)

Cytochrome *bo*_3_ (c*bo*_3_) proton translocation was tracked in a method similar to that in Hards *et al*.^1^. Briefly, proteoliposomes (0.2 mg) consisting of 2% c*bo*_3_/mass *E. coli* polar lipids doped with 1% mass ubiquinone-10 (UQ_10_) per ml were pre-warmed to 37 °C for 15 min in 20 mM MOPS, 30 mM Na_2_SO_4_, pH 7.4, 1 mM DTT, and 1 μM ACMA ± 1μM valinomycin with vigorous stirring (800 rpm). Quenching was initiated by the addition of 2.5 μM ubiquinone-1 (UQ_1_) in ethanol and reversed as indicated in text. Ethanol controls had no effect on ACMA quenching.

### Electrochemical assays

Electrochemical experiments in tethered bilayer lipid membranes and proteoliposomes were prepared as described elsewhere^8^. All the experiments were carried out with ultra-flat template stripped gold (TSG) working electrodes. 150 nm of 99.99% gold (Goodfellows) was evaporated on silicon wafers using a Telemescal evaporator at < 2×10^−6^ mbar. 1.2 cm^2^ glass slides were glued to the gold layer with Epo-Tek 377 and cured for 2 h at 120°C. The TSG surface was exposed by detaching the glass slides from the silicon wafers before use. The formation of the self-assembled monolayers (SAMs) containing the cholesterol “tether” and the formation of the SSM onto the electrode were performed as described previously^8^. SAMs were formed by incubating a freshly exposed TSG slide in 0.11 mM EO3-cholesteryl and 0.89 mM 6-mercaptohexanol (6MH) in propanol for 16 h. Where vesicles/proteoliposomes were used as described previously^9^, SAMs were made with 1 mM 6MH alone. After incubation, the excess thiol was gently washed away with isopropanol and methanol, and the electrodes were then dried in a stream of N2. For bilayer SAMs this procedure results in an approximate 60%/40% EO3-cholesteryl/6-mercaptohexanol area ratio on the surface as confirmed by impedance spectroscopy before each experiment. For SAMs prepared for vesicle/proteoliposome experiments, the surface was coated in 100% 6MH. To form tethered lipid membranes (tBLMs), vesicles or proteoliposomes were added to the SAM surface at a final concentration of 0.5 mg/mL in the presence of 10 mM CaCl_2_ and incubated for 1h until a capacitance drop to less than 1.2 μF/cm^2^ was observed. The surface was then rinsed three times with water, then buffer containing 0.5 mM EDTA to remove any traces of calcium ions in the cell. In vesicle/proteoliposome experiments, either vesicles or proteoliposomes were added to the 100% 6MH SAM surface at a final concentration of 0.5 mg/mL and incubated for 30min. Finally, the SAM-modified or vesicle/proteoliposome-modified electrodes were rinsed three times with buffer and used in the electrochemistry experiments. Care was taken to keep the electrodes immersed in an aqueous environment at all times during rinsing. The time window for measurable proton leak is 25 s from quinone reduction on the forward scan to the start of the reverse scan.

### Quantification of detergents by LC-MS/MS

Protein extracts and detergent standards were analysed using a UPLC BEH (1.0 × 100 mm, 1.7 μm, Waters, Acquity) separation phase coupled online to a ESI-Q-TOF mass spectrometer (Q-TOF Premier, Waters Micromass, Manchester, UK) operated in ESI+ mode, alternating full scan and selected reaction monitoring (SRM) modes. After sample injection, a constant flow at 50 μL/min of a solvent consisting of 15% B was kept over 3 minutes, followed by a linear gradient to a solvent composition of 95 % B over 16 minutes. Finally, the flow was kept constant at 95% B over additional 4 minutes until the back-equilibration to solvent start conditions. The flow rate of 50 μL/min was maintained using an Acquity Ultra Performance LC pump system (Waters, Milford, United States). Buffer A consisted of 50 mM ammonium acetate in LC-MS grade water and buffer B consisted of 5 % mobile phase A in LC-MS grade acetonitrile. During all analysis runs, samples were cooled at 10°C and the analytical separation column was maintained at 25°C. The full scan was acquired over the mass range of 150 – 1250 m/z at 1 sec scan times, and the selected reaction monitoring (SRM) modes isolated mass windows at 528.3 m/z, for dodecyl-D-maltoside (DDM) and 1005.6 m/z for Lauryl maltose neopentyl glycol (LMNG), over 1 sec scan times each. A collision energy (CID) of 9eV was used for selected reaction monitoring. The Q-TOF mass spectrometer was calibrated using Glu1-Fibrinopeptide B (human, Sigma-Aldrich) before analysis.

External calibration for detergents was performing by injecting a dilution series of every detergent (DDM: 3.82^−3^ to 5.0^−5^ μmol on column injection, LMNG: 1.6^−4^ to 4.0^−5^ μmol on column injection) to the LC-ESI-Q-TOF analysis system. All standards and samples were analyzed in duplicates and summed peak area counts of the major fragment of the respective selected reaction monitoring trace were used to establish a calibration curve and to further determine the concentration of the detergents in the protein extracts. Data analysis was performed manually using MassLynx 4.1, and determination of calibration curves and calculation of detergent concentrations in samples was performed using Microsoft Excel (for calibration curves see supplemental information material).

### Negative stain EM of liposome-reintegrated *B. taurus* F-type ATP synthase

A 2.5 μl aliquot was applied onto freshly glow-discharged, carbon-coated 400 mesh copper grids (Veco). After brief blotting (Whatman #1), the samples were stained by using a 2% uranyl acetate solution and air-dried. Images were taken with a JEM1010 transmission electron microscope (JEOL) equipped with a 4 x 4 K Tietz CMOS TemCamF416 (TVIPS, Gauting, Germany) at 100 kV.

### Cryogenic electron tomography of liposome-reintegrated *B. taurus* F-type ATP synthase

QUANTIFOIL (R0.6/1, Mo) grids with gold colloidal markers on the carbon films were used for cryo-EM. The grids were coated with poly-L-lysine (BBI Solutions) for 10 min after 1 min glow-discharging and then 15 nm colloidal golds (Sigma-Aldrich) were applied for 10 min. The solution was removed from the grids with filter paper and then the grids were washed by distilled water. The grids were coated with a 10- to 20-nm-thick layer of carbon after drying. 3 μl of sample was applied onto the grid and blotted for 6 s with a blot force of 10 at 4 °C and 100% humidity using a Vitrobot (Thermo Fisher) and then flash-frozen by plunging into liquid ethane. The samples were observed with a Titan Krios (Thermo Fisher) operating at 300 kV and equipped with a Falcon II detector at a direct magnification of 22.5K. The images were taken from 0° to +70° and then from −2° to −70° with 2° steps. The defocus values were about 5 μm and the electron dose for each exposure was 1.6 e^−^/Å^2^ with a total dose of less than 120 e^−^/Å^2^. Tilt series were aligned using the gold fiducials and back-projected to generate tomographic volumes using the IMOD package^11^ with a pixel size of 5.7 Å.

All studies reported here studies have complied with all relevant ethical regulations for animal testing and research.

### Data Availability

Data supporting the findings of this manuscript are available from the corresponding authors upon reasonable request. Video files of cryo-tomography are available as supplementary information.

## Notes

### Competing Interest Statement

The authors have declared no competing interest.

