## Supplementary data for "Autoinsertion (LAiR): rapid functional reconstitution of integral membrane proteins into lipid bilayers"

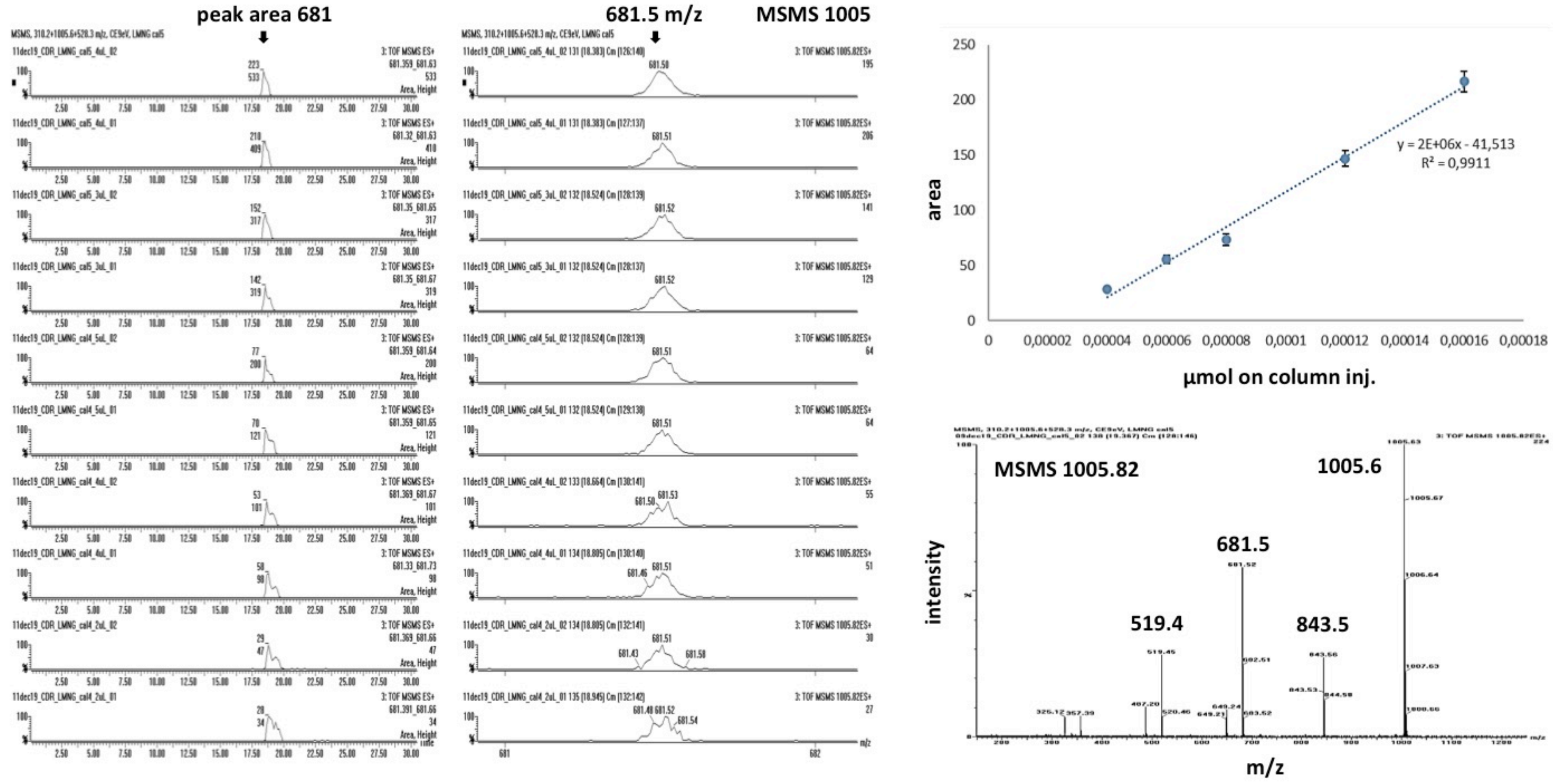


**Supplemental Figure 1:** SRM analysis of a lauryl maltose neopentyl glycol (LMNG) standard dilution series (from $1.6^{-4}$ to $4.0^{-5}$ µmol on column in duplicates, 1005.6 m/z) including calibration curve (upper right graph, using the area of major fragment 681.5 m/z) and an annotated fragmentation spectrum (lower right spectrum).

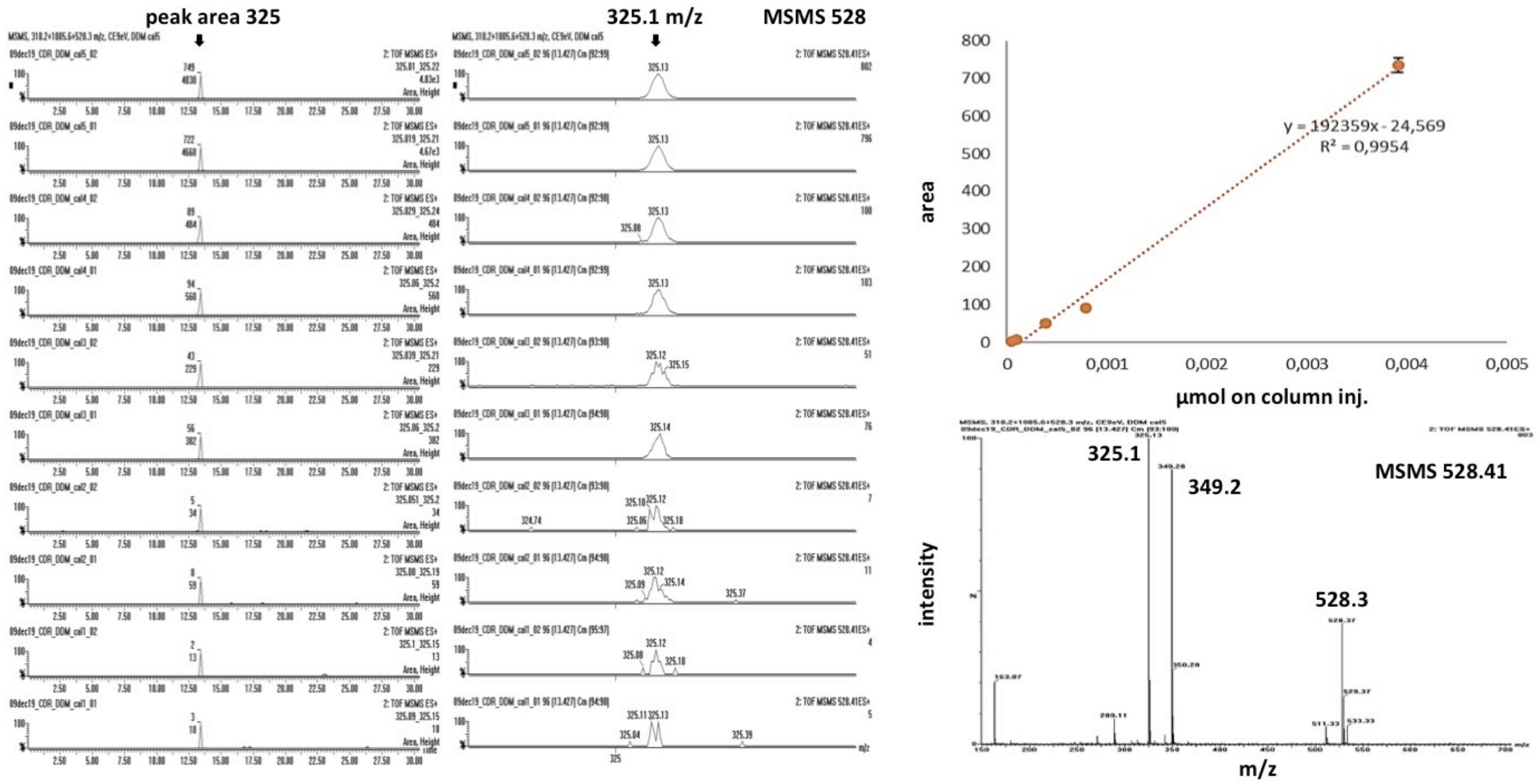


**Supplemental Figure 2:** SRM analysis of a β-dodecyl-maltoside (DDM) standard dilution series (from $3.82^{-3}$ to $5.0^{-5}$ µmol on column in duplicates, 528.4 m/z) including calibration curve (upper right graph, using the area of major fragment 325 m/z) and an annotated fragmentation spectrum (lower right spectrum).
